# Opposing directions of stage-specific body length change in a close relative of *C. elegans*

**DOI:** 10.1101/2020.06.23.168039

**Authors:** Eric W. Hammerschmith, Gavin C. Woodruff, Patrick C. Phillips

**Affiliations:** Institute of Ecology and Evolution, University of Oregon, Eugene, Oregon, USA; Princeton Neuroscience Institute, Princeton University, Princeton, New Jersey, USA

**Keywords:** Body size, phenotypic plasticity, ontogenetic niche, *C. elegans*

## Abstract

**Background:** Body size is a fundamental organismal trait. However, as body size and ecological contexts change across developmental time, evolutionary divergence may cause unexpected patterns of body size diversity among developmental stages. This may be particularly evident in polyphenic developmental stages specialized for dispersal. The dauer larva is such a stage in nematodes, and *Caenorhabditis* species disperse by traveling on invertebrate carriers. Here, we describe the morphology of the dispersal dauer larva of the nematode *Caenorhabditis inopinata*, whose adults can grow to be nearly twice as long as its close relative, the model organism *C. elegans*.

**Results:** We find that the *C. inopinata* dauer larva is shorter and fatter than those of its close relatives *C. elegans, C. briggsae*, and *C. tropicalis*, despite its much longer adult stage. Additionally, many *C. inopinata* dauer larvae were ensheathed, an apparent novelty in this lineage reminiscent of the infective juveniles of parasitic nematodes. We also found abundant variation in dauer formation frequency among twenty-four wild isolates of *C. inopinata*, with many strains unable to produce dauer larvae under laboratory conditions.

**Conclusion:** Most *Caenorhabditis* species thrive on rotting plants and disperse on snails, slugs, or isopods (among others) whereas *C. inopinata* is ecologically divergent and thrives in fresh *Ficus septica* figs and disperses on their pollinating wasps. These wasps are at least an order of magnitude smaller in length than the vectors of other *Caenorhabditis* species. While there is some unknown factor of the fig environment that promotes elongated body size in *C. inopinata* adults, the smaller size of its fig wasp carrier may be driving the reduced body length of its dauer larva. Thus ecological divergence across multiple developmental stages can promote unexpected and opposing changes in body size within a single species.

## Background

One obvious fact of the life cycles of most organisms is that they typically get larger as they develop. Indeed, most animals span at least an order of magnitude in body size within a single generation through the course of development [1, 2]. However, a frequently neglected consequence of this is that the ecological niche of an organism can change drastically within a single individual throughout development. Notable examples of such size-structured ecological niches (i.e., ontogenetic niches [3]) include animals exhibiting metamorphosis in development. Ceratophryidae frogs have larval forms that eat small crustaceans and diatoms yet grow into adult forms that eat larger arthropods, gastropods, and small vertebrates [4]. Silkworms eat mulberry leaves as larvae yet develop into short-lived, non-feeding adults specialized for reproduction [5, 6]. Such stage-specific ecology is not restricted to animals with dramatic metamorphic development. Mammals usually begin postembryonic life consuming milk before maturing into adults that consume plants, other animals, or both [1]. Many species of fish also exhibit stage-specific resource use, and largemouth bass eats planktonic crustaceans, crawfish, and cyprinid fish as it grows [3, 7].

The ontogenetic niche of an organism is not limited by resource use. Organisms often have specialized developmental stages for specific life history strategies, and such partitioning is frequently used for dispersal. Locusts have a complex polyphenism in overcrowding conditions that transform solitary morphs into gregarious morphs that constitute famine-inducing swarms [8]. Aphids also have a polyphenism specialized for dispersal where winged morphs arise in harsh conditions; dimorphic aphids also are easily distinguished by thorax size differences [9]. The seeds of plants and spores of fungi are small stress-resistant propagules that frequently harbor traits that aid in dispersal such as wings or barbs (in the case of seeds) [10, 11]. Thus life history strategies can also influence stage-specific traits including size.

As adults, the free-living bacterivorous nematodes *Caenorhabditis elegans* and *C. inopinata* live in substantially different habitats and differ substantially in size. In crowded, low food, high temperature, or otherwise stressful conditions, *C. elegans* is known to develop into a dispersal-specialized juvenile phase known as the dauer larva [12]. The dauer larva is a long-lived, desiccation-resistant, and stress-resistant dispersal stage. In its natural context, *C. elegans* dauer larvae travel to new resources on invertebrate carriers such as snails, slugs, isopods, and myriapods [13]. The dauer exhibits a stage-specific behavior in nictation, wherein the animal climbs a substrate and waves its head in the air to promote invertebrate-mediated dispersal [14]. Once it has traveled to a new rotting plant resource patch, the dauer larva will disembark and directly develop into a reproductive stage animal [13]. As the dauer larva is a specialized L3 stage, it has a distinct morphology from reproductive phase animals [12]. While a great deal is known about the *C. elegans* dauer stage, the dauer stage from *C. inopinata* has never before been described. Do the substantial differences observed in the adult species of these close relatives persist throughout this developmental stage as well?

Here, we describe the dauer larva of *Caenorhabditis inopinata*. While *C. inopinata* adults grow twice as long and *C. elegans*, they also develop nearly twice as slowly as *C. elegans* [15-17]. Additionally, instead of thriving on rotting plants and dispersing on large invertebrates, *C. inopinata* lives in fresh *Ficus septica* figs and disperses on fig wasp pollinators [15, 18]. Here we show that, rather than echoing the size differences observed in the adults, *C. inopinata* dauers are shorter and fatter than *C. elegans* dauers. In addition, *C. inopinata* dauers also exhibit a novelty in ensheathment that resembles the infective larvae of parasitic nematodes. Thus the evolution of body size need not be in the same direction across all developmental stages, and stage-specific ecological contexts can drive opposing directions of body size change.

## Results

*C. inopinata* adults are much longer than those of *C. elegans* (Figure 1A; [15, 16]). However, as *C. inopinata* disperses on fig wasp vectors that are millimeters long, we were curious about the extent of body size change in its dispersal dauer larva. In *C. elegans*, dauers are typically isolated via SDS exposure [19], which kills non-dauer stages that can feed and lack a buccal plug. This approach has also been shown to isolate dauer larvae in *C. briggsae* [20]. We also successfully used this approach to isolate and characterize the morphology of *C. inopinata* dauer larvae (Fig 1. B-D). The vast majority of *C. inopinata* animals that survived SDS exposure had a characteristic buccal plug (Fig. 1C; 98%, 45/46 worms), revealing this is a sound method of recovering animals of the appropriate stage. Surprisingly, despite the elongation of *C. inopinata* adults compared to *C. elegans* (Fig. 1A), *C. inopinata* dauers are shorter (*C. inopinata*: mean = 430 microns, SD = 45 microns; *C. elegans*: mean= 456 microns, SD = 65 microns; Wilcoxon rank sum test *p* = 0.0092, W = 4337) and fatter (*C. inopinata*: mean = 27 microns, SD = 4.1 microns; *C. elegans*: mean= 18 microns, SD = 3.5 microns; Wilcoxon rank sum test *p* < 2.2 × 10^−16^, W = 10530) than its close relatives (Fig. 1-2). This leads to the dauer larva occupying different regions of morphological space in *C. elegans* and *C. inopinata* when compared with other developmental stages (Figure 2A). And while *C. elegans, C. briggsae*, and *C. tropicalis* differ in dauer length (Figure 2B; Tukey test *p* = 1.4 × 10^−11^), they have negligible differences in width (*C. tropicalis* is on average one micron wider than *C. elegans*; Tukey test *p* = 0.020), revealing the *C. inopinata* dauer to be an outlier with a length:width ratio that is 40% smaller than its close relatives (Figure 2B). Thus *C. inopinata* has a divergent dauer stage morphology that is surprisingly inconsistent with its elongated adult stage. Additionally, a notable fraction of *C. inopinata* dauer larvae retained their cuticle from the previous molt (46%, 21/46 worms). Reminiscent of the ensheathed infective larvae of parasitic nematodes [21, 22], this was never observed in *C. elegans* dauers (n=47 worms).

**Figure 1.**
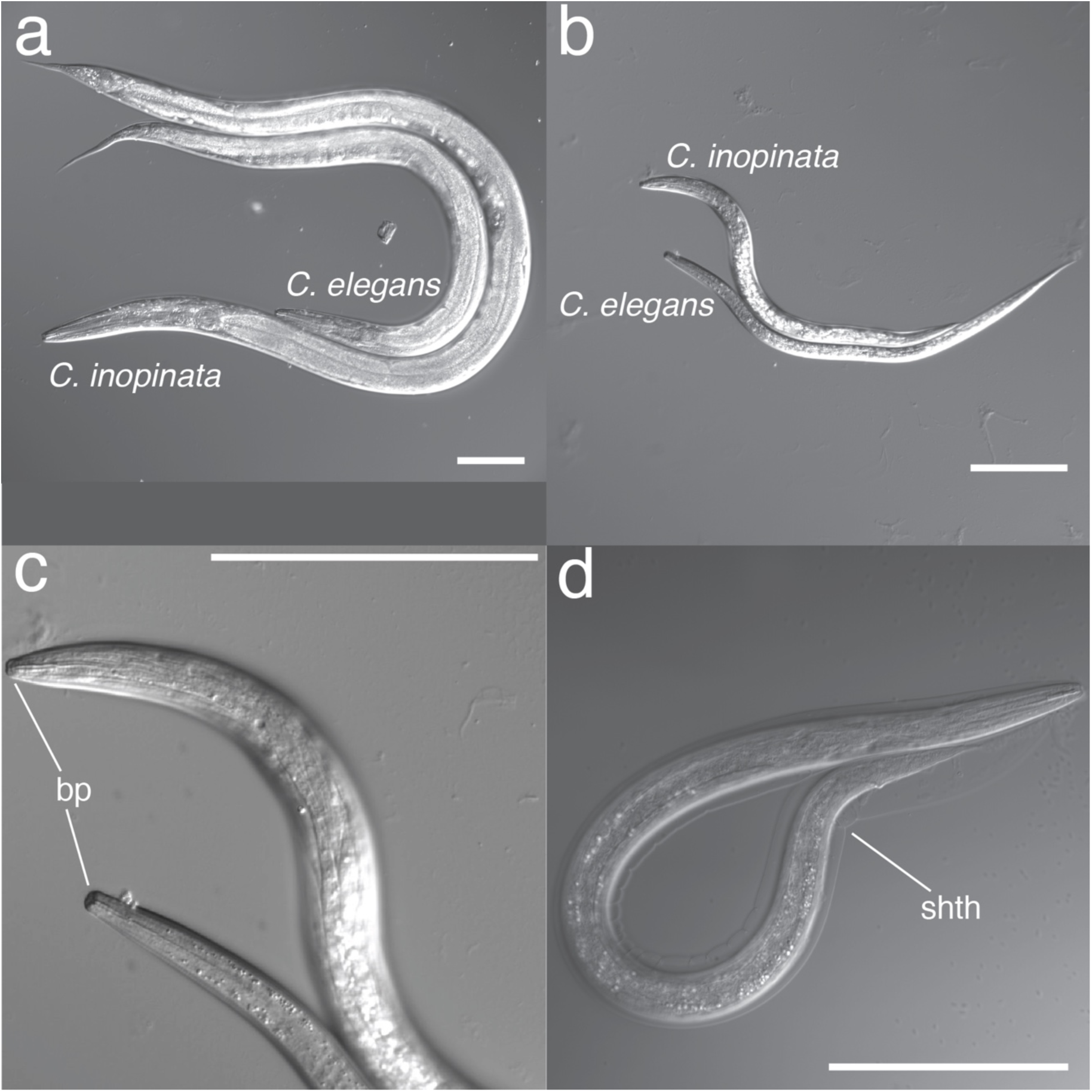
The *C. inopinata* dauer is morphologically distinct. (a) *C. elegans* adult hermaphrodite with *C. inopinata* adult female. (b) *C. elegans* and *C. inopinata* dauer larvae. (c) Enlargement of panel (b) showing the buccal plug. Bp, buccal plug. (d) An ensheathed *C. inopinata* dauer larva. Shth, sheath. All scale bars represent 100 microns.

**Figure 2.**
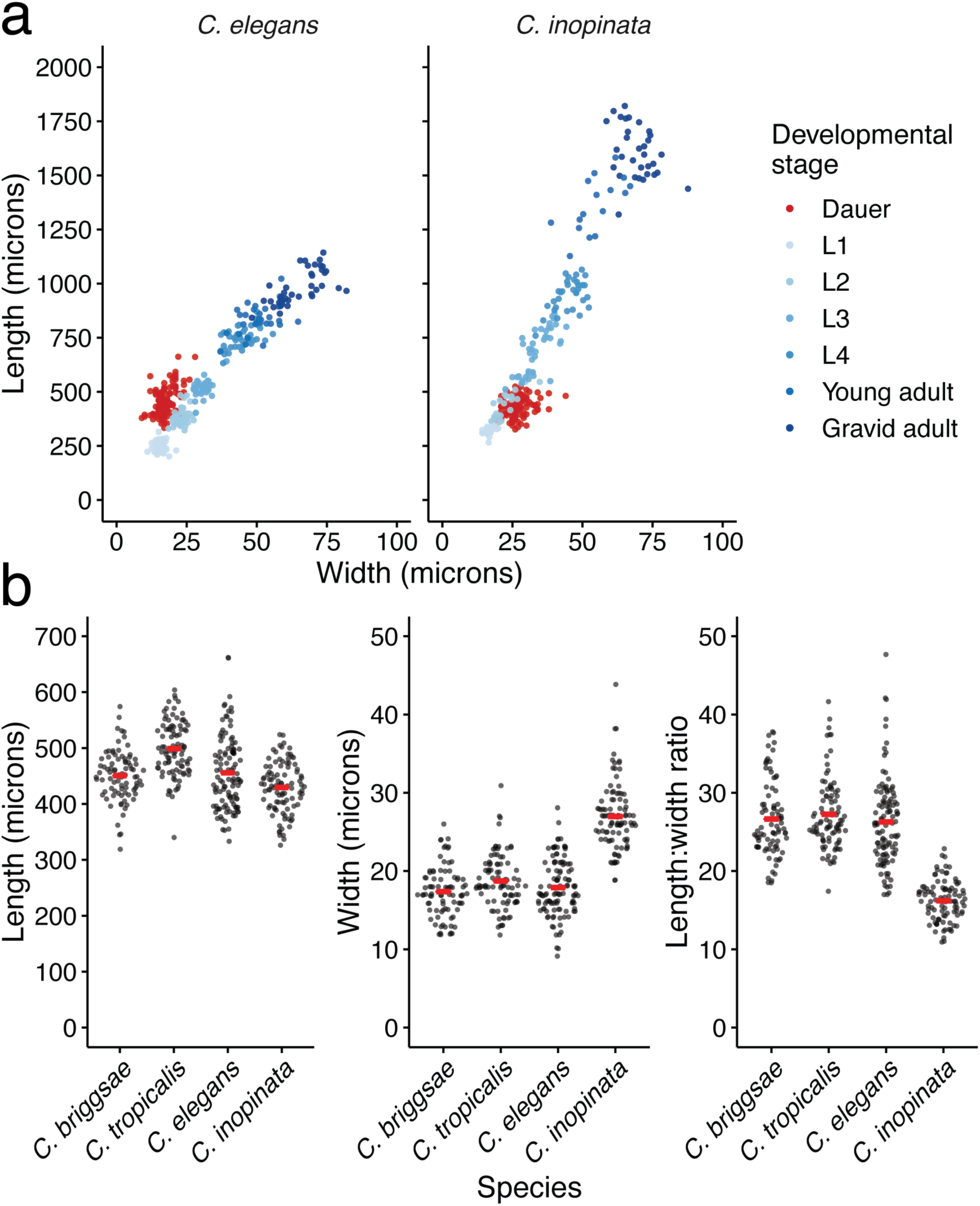
Quantification of dauer size. (a) Scatterplots revealing the length and width of various developmental stages. Data for non-dauer stages are from Woodruff et al. 2018. Left panel, *C. elegans*; Right panel, *C. inopinata*. Non-dauer stages N_worms_= 16-103; *C. elegans* dauers, N_worms_= 165; *C. inopinata* dauers, N_worms_= 132. (b) Sina plots (strip charts with points taking the contours of a violin plot) illustrating the distributions of dauer length (left panel), dauer width (middle panel), and the dauer length:width ratio (right panel) of four *Caenorthabditis* species. Each dot represents the observation of one worm. Red bars represent averages. Sina plots take *C. elegans*, N_worms_ = 113; *C. inopinata*, N_worms_ = 97; *C. briggsae*, N_worms_ = 85; *C. tropicalis*, N_worms_ = 93.

Anecdotally, it was clear that we were recovering low numbers of *C. inopinata* dauers in our SDS treatments. As variation in dauer formation frequency has long been noted in *C. elegans* [23-25], we attempted to isolate dauers from a number of *C. inopinata* wild strains from Okinawan *Ficus septica* figs (Figure 3; Supplemental Table Sheets 1-2; Supplemental Figure 1). As observed in *C. elegans*, there is variation in dauer formation frequency in *C. inopinata* among 29 lines (Figure 3; range = 0-2%, mean = 0.2%, sd= 0.4). However, most lines generate dauers at a low frequency, and many lines never produced dauers (11 lines; Figure 3). Furthermore, there appears to be no relationship between island of origin and dauer formation frequency (Figure 3; Wilcoxon rank sum test *p* = 0.47, W = 4068). This is in contrast to other *Caenorhabditis* species, where many lines are able to produce abundant dauers in starvation conditions (Supplemental Figure 2, [20, 23-25]). Thus the ecological divergence of *C. inopinata* may have impacted the dauer entry and exit decisions in this species, leading to its low propensity to promote dauer formation in laboratory conditions.

**Figure 3.**
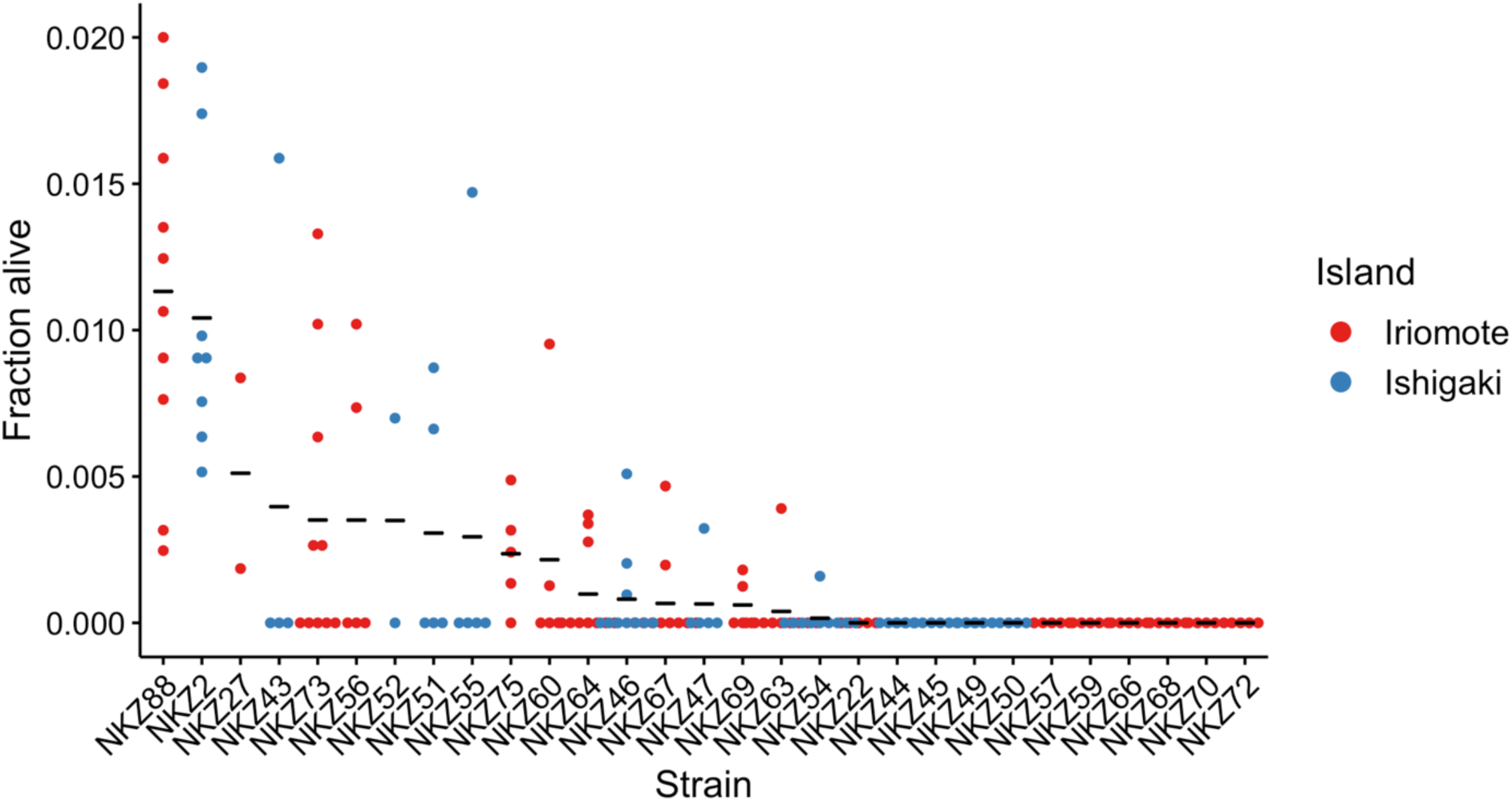
Variation in dauer formation frequency in *C. inopinata.* Twenty-five *C. inopinata* strains are shown on the x-axis; all are wild isolates with the exception of NKZ43, which is a twenty-generation inbred line derived from NKZ2. Each point represents one observation of dauer formation frequency, the fraction of a starved population of animals surviving SDS treatment (N observations per strain = 2-10 ; N worms per observation = 17- 1557, average = 450, median = 397). Black horizontal lines represent the average. Strains are colored by Okinawan island of origin (Supplemental_tables.xls Sheet 1).

## Discussion

The partitioning of life history strategies among developmental stages is widespread. Among the most speciose groups of animals, holometabolous insects have just this kind of life cycle: larvae are specialized for growth while adults are suited for reproduction and dispersal [26]. This sorting leads to dramatic morphological and ecological differentiation among stages, with adults being nearly unrecognizable compared to their younger forms. Thus the allotment of such strategies can drive marked diversity across development within a single organism. And although many animals do not undergo striking metamorphosis throughout their development, diapause and dispersal are pervasive approaches for responding to environmental change and for promoting reproductive assurance [27]. These are frequently tied to specialized stages and often require phenotypic plasticity to mediate entrance and exit from such forms at the appropriate time [28]. And as this is usually accompanied by morphological and physiological adaptations suited for these strategies, the partitioning of life history strategies across development provides ample opportunity for the generation of variation both within and between individuals.

The nematode dauer larva represents such a stage: it is a stress-resistant, long-lived alternative developmental stage specialized for dispersal whose development is induced by adverse environments. Here, we described the broad morphological characteristics of the *C. inopinata* dauer larva and found that it is shorter and fatter than the dauers of other *Caenorhabditis* species, including *C. elegans* (Fig. 1-2). This finding is unexpected as *C. inopinata* is easily distinguished from its close relatives by its exceptionally elongated adult body size [15, 16]. Indeed, *C. inopinata* has been reported to grow as much as nearly twice [16] to three times [15] longer than *C. elegans*. Thus its comparatively shorter dauer larva is seemingly incongruent with its longer adult form. However, this observation can be clarified by situating *Caenorhabditis* dauer larvae in their ecological contexts.

Most *Caenorhabditis* species thrive by consuming the microbes in rotting plant material such as stems, fruit, and flowers [13, 29]. In favorable environmental conditions, animals directly develop into reproductively mature adults. In poor conditions, animals develop into the alternative dauer larval stage [12]. Dauer larvae harbor a number of stage-specific traits that promote dispersal including nictation, wherein animals climb a substrate and wave their heads in the air [14], and experimental evidence suggests this behavior aids in phoresis on invertebrate carriers [30]. *Caenorhabditis* dauer larvae have been frequently observed on such vectors, and they presumably carry dauers to new rotting plant resources where they disembark and mature. *C. elegans* has been observed on snails (*Helix, Oxychilus*, and *Pomatias* species [31, 32]), slugs (*Deroceras, Arion*, and *Ambigolimax* species [31, 33, 34]), isopods (*Porcellio, Oniscus*, and *Armadillidium* species [33, 35, 36]), millipedes (*Glomeris* species [32]), and centipedes [33, 37]. *C. briggsae* and *C. remanei* have also been observed on similar vectors (gastropods, isopods, and myriapods [13, 34, 37-39]. Thus some *Caenorhabditis* species can disperse on a diverse array of invertebrate carriers than span ∼10-65 mm in body length (Fig. 4; [40, 41]). *C. inopinata*, on the other hand, thrives in an entirely different ecological context, the lumen of fresh *Ficus septica* figs [15, 18]. Furthermore, instead of dispersing on a variety of invertebrate carriers, *C. inopinata* primarily travels on *Ceratosolen* pollinating fig wasps [15, 18]. Although *C. inopinata* has been observed on the parasitic fig wasp *Phylotrypesis* [15], it appears to preferentially embark on *Ceratosolen* pollinators [15, 18]. Thus *C. inopinata* is notable in its degree of host vector specialization. Furthermore, fig wasp vectors are up to two orders of magnitude smaller in length than the vectors of *C. elegans* and other rotting plant-associated *Caenorhabditis* species (Fig. 4; [42]). Thus, although there must be some (as yet unknown) factor of the fig environment driving increased adult body length in *C. inopinata* adults, the divergent morphology of its dauer larvae can likely be at least partly explained by its need to disperse on much smaller vectors (Fig. 4). In this way, the divergent ecological contexts of different developmental stages can drive morphological divergence of such stages in opposing directions.

**Figure 4.**
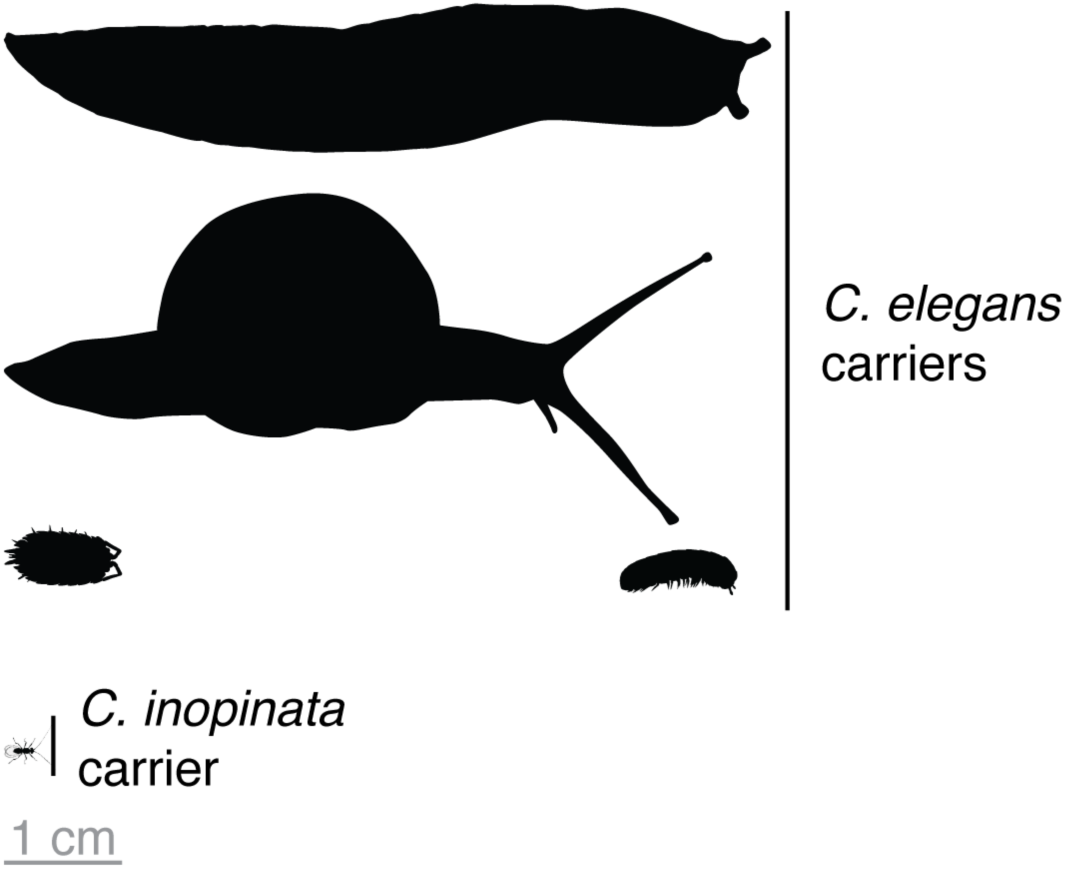
Variation in the size of *Caenorhabditis* vectors. Silhouettes of invertebrates are to scale. *C. elegans* has been observed traveling on *Arion* slugs [33] (∼65 mm long [41]), *Helix* snails [31] (∼60 mm long [56]), *Porcellio* isopods [33, 35, 36] (∼10 mm long [40]), and *Glomeris* millipedes [32] (∼10 mm long [57]). *C. inopinata* disperses on *Ceratoslen bisculatus* fig wasps [15, 18], which are much smaller in size (∼1.7 mm long [42]). All silhouettes were retrieved from PhyloPic (http://phylopic.org/) and designed by Birgit Lang (isopod), Gareth Monger (gastropods), Thomas Hegna (millipede), and Kamil S. Jaron (wasp). Adult *Caenorhabditis* nematodes are roughly the size of the crossbar in the lowercase “t,” and dauers are roughly the size of a period. A version of this figure including nematodes to scale is in the supplemental material (Supplemental Figure 3).

Such ecological divergence in host vectors may also explain the novel ensheathment trait observed in *C. inopinata* dauer larvae (Fig. 1D). The retention of the L2 cuticle in diapause larvae, or ensheathment, is widespread in nematodes [21]. Often associated with the infective larvae of parasitic species, ensheathment is thought to aid in desiccation resistance [43], predation resistance [44], and freezing tolerance [45]. Furthermore, in many parasitic nematodes, passage through an ensheathed infective (i.e., dauer) larval stage is essential for progression through the life cycle [21, 46]. At the same time, it is important to note that ensheathment is not restricted to parasites and has been observed in free-living nematodes [47]. To the best of our knowledge, ensheathment has never been observed before in *Caenorhabditis* nematodes. This includes the vector specialist *C. drosophilae*, whose dauer larva is also associated with a comparatively small host (*Drosophila nigirospiracula*; thorax length ∼1.3 mm [48]) but is not ensheathed [49]. Thus it is possible that vector size alone is not driving the evolution of this trait in *C. inopinata*. Dissections of *Ceratosolen* fig wasps have revealed the presence of *C. inopinata* dauer-like larvae in wasp hemolymph [15]; thus ensheathment may be important for dauer survival in this novel environment. However, presumably unsheathed *Caenorhabditis* species have been observed in the guts of snails and slugs [33, 38], so ensheathment is possibly not needed for the survival of internal passage in invertebrate carriers *per se*. Additionally, although *C. inopinata*’s relationship with fig wasps is thought to be phoretic (like with other *Caenorhabditis* species and their invertebrate carriers), it remains possible that they negatively impact fig wasp fitness. Although direct fig wasp parasitism seems unlikely as *C. inopinata* is bacterivorous and culturable in the lab without fig wasps [15-17], *C. inopinata* may be necromenic or entomopathogenic. Entomopathogenic *Steinernema* species have ensheathed larvae and carry the pathogenic bacteria that kills their slug hosts and allows them to thrive [50]. Furthermore, the dauer larvae of fig wasp-associated *Parasitodiplogaster* species have also been observed to be ensheathed [51]. This could be further evidence that ensheathment may be needed for successful transit on fig wasps in particular. However, fig-associated *Schistonchus* nematodes often do not have apparent dauer larvae (with or without ensheathment) as the infective stage is usually the pre-reproductive female [52]. Regardless, as *C. inopinata* has gained a trait commonly associated with animal parasitism, this affords the opportunity of using the *Caenorhabditis* model system genetics toolkit to potentially understand the evolution of parasitism-related traits.

Variation in dauer formation frequency was also observed in *C. inopinata* (Fig. 3). In *C. elegans*, there is substantial variation in this trait [23-25], and it has a complex genetic basis, with at least thirty-six loci contributing to most of the variance in dauer formation frequency [25]. Additionally, most isolates produced small numbers of dauer larvae, with many producing none at all (Fig. 3). As it is likely that fig and/or fig wasp cues influence the dauer decision in natural conditions [18], it is possible that their absence in laboratory culture promotes the low dauer formation frequencies observed in *C. inopinata*. Future studies will be needed to disentangle the roles of dauer entry, exit, and their interactions with figs and fig wasps in the variation of this trait. Furthermore, we observed no clear relationship between island of origin and dauer formation frequency in these lines; this is consistent with preliminary population genomic data that reveal little genetic differentiation between Okinawan island populations (Woodruff et al., manuscript in prep.). Regardless, future quantitative genetic studies using these isolates will prove invaluable for understanding the genetic basis of dauer formation and other quantitative traits in *C. inopinata*.

## Conclusions

Here, we showed that the *C. inopinata* dauer larva has decreased length despite its elongated adult body size compared to *C. elegans*. This is likely due to the divergent ecological contexts of these developmental stages that are specialized for dispersal or reproduction. Fig wasps are the dispersal vector of *C. inopinata* that is at least an order of magnitude smaller than those used by *C. elegans*, while figs have potentially released some selective constraint that allows larger adult body sizes compared to the rotting plant environments of *C. elegans*. Thus polyphenisms and specialized developmental stages reveal surprising flexibility in the direction and magnitude of body size evolution in a single species.

## Methods

### Strains and maintenance

Animals were grown on modified Nematode Growth Media with *Escherichia coli* strain OP50-1 for food as in [17]. All animals were raised at 25°C. Strains *C. inopinata* NKZ2 (also known as NK74SC [15, 16]), *C. elegans* N2, *C. briggsae* HK104, and *C. tropicalis* NIC122 were used for dauer morphology observations. For characterizing variation in dauer formation frequency, multiple strains of *C. elegans* (ED3040, CB4856, JU1088, JU775, MY16), *C. briggsae* (JU1348, ED3091, HK104), and *C. tropicalis* (NIC122, QG131) were used.

We report multiple new wild isolates of *C. inopinata* used for characterizing dauer formation variation. *C. inopinata* strains NKZ22 and NKZ27 were isolated from fresh *F. septica* figs from the island of Iriomote, Okinawa, Japan in May 2014 (Supplemental Figure 1; supplemental_tables.xls, Sheet 1). *C. inopinata* strains NKZ44, NKZ45, NKZ46, NKZ47, NKZ49, NKZ50, NKZ51, NKZ52, NKZ54, NKZ55, NKZ56, NKZ57, NKZ59, NKZ60, NKZ63, NKZ64, NKZ66, NKZ67, NKZ68, NKZ69, NKZ70, NKZ72, NKZ73, NKZ75, and NKZ88 were isolated from fresh *F. septica* figs from the islands of Iriomote and Ishigaki, Okinawa, Japan in May 2015 (Supplemental Figure 1; supplemental_tables.xls, Sheet 1). *C. inopinata* strains PX723 and PX724 were isolated from fresh *F. septica* figs from Taipei, Taiwan in August 2019 (Supplemental Figure 1; supplemental_tables.xls, Sheet 1). Wild isolates were established as in [16]. Briefly, figs were placed in a petri dish filled with water or M9 buffer. Figs were then cut into four pieces; worms subsequently found in suspension were placed on to NGM plates to establish wild isolates. Morphology and association with *F. septica* strongly suggest these wild isolates are the same species as *C. inopinata*. Molecular barcoding with the ITS2 (ribosomal rDNA internal transcribed spacer-2) sequence and/or mating tests were performed in a few of these lines, confirming their species identity (supplemental_tables.xls, Sheet 2; [53]). As mating tests have not been performed in a number of lines, there exists the unlikely possibility these particular lines are cryptic species distinct from *C. inopinata. C. inopinata* strain NKZ43 is an inbred line of *C. inopinata* made through 20 generations of sib-pair inbreeding (derived from the genome-sequenced strain NKZ35 [15]).

### Dauer isolation

Dauer larvae were isolated 10-14 day old starved cultures incubated at 25°C. Animals were washed off of plates in M9 buffer and then incubated in 1% SDS for a half hour. Animals were then washed four times in M9 buffer and plated. Live worms were then used for subsequent observations.

### Microscopy and measurements

Animals were mounted on agar pads and imaged on a dissecting microscope. Animal length and width was measured using the ImageJ software [54]; curved lines were addressed with the “segmented line tool” as in [55]). All measures of reproductive, non-dauer stages were retrieved from the data in [16]. The fraction of animals surviving SDS treatment was determined by counting the number of live and dead worms from a sample of a given suspension both before and after SDS exposure. All statistics were performed in the R language, and all data and code associated with this study have been deposited on Github (https://github.com/gcwoodruff/dauer_2020).

## List of abbreviations

L3: third larval stage.
SDS: Sodium dodecyl sulfate.

## Declarations

### Ethics approval and consent to participate

Not applicable. *Consent for publication* Not applicable.

### Availability of data and material

All data and code not included as Additional Files have been posted to Github (https://github.com/gcwoodruff/dauer_2020).

### Authors’ contributions

GCW, PCP, and EH designed the experiments; EH and GCW performed the experiments; EH and GCW analyzed the results; EH, GCW, and PCP wrote the paper.

## Acknowledgements

We would like to thank John Wang and Natsumi Kanzaki for their invaluable assistance in the isolation of *C. inopinata* strains.

## Additional Files

Additional File 1. Supplemental Figures (Supplemental_figures.doc). Additional File 2. Supplemental Tables (Supplemental_tables.xls).

## Supplemental Figures

Supplemental Figure 1. Localities of *C. inopinata* strains isolated from Okinawa (islands of Iriomote and Ishigaki) and Taiwan.

Supplemental Figure 2. *C. inopinata* produces few dauer larvae compared to its close relatives.

