## supplemental_figures for "Opposing directions of stage-specific body length change in a close relative of *C. elegans*"

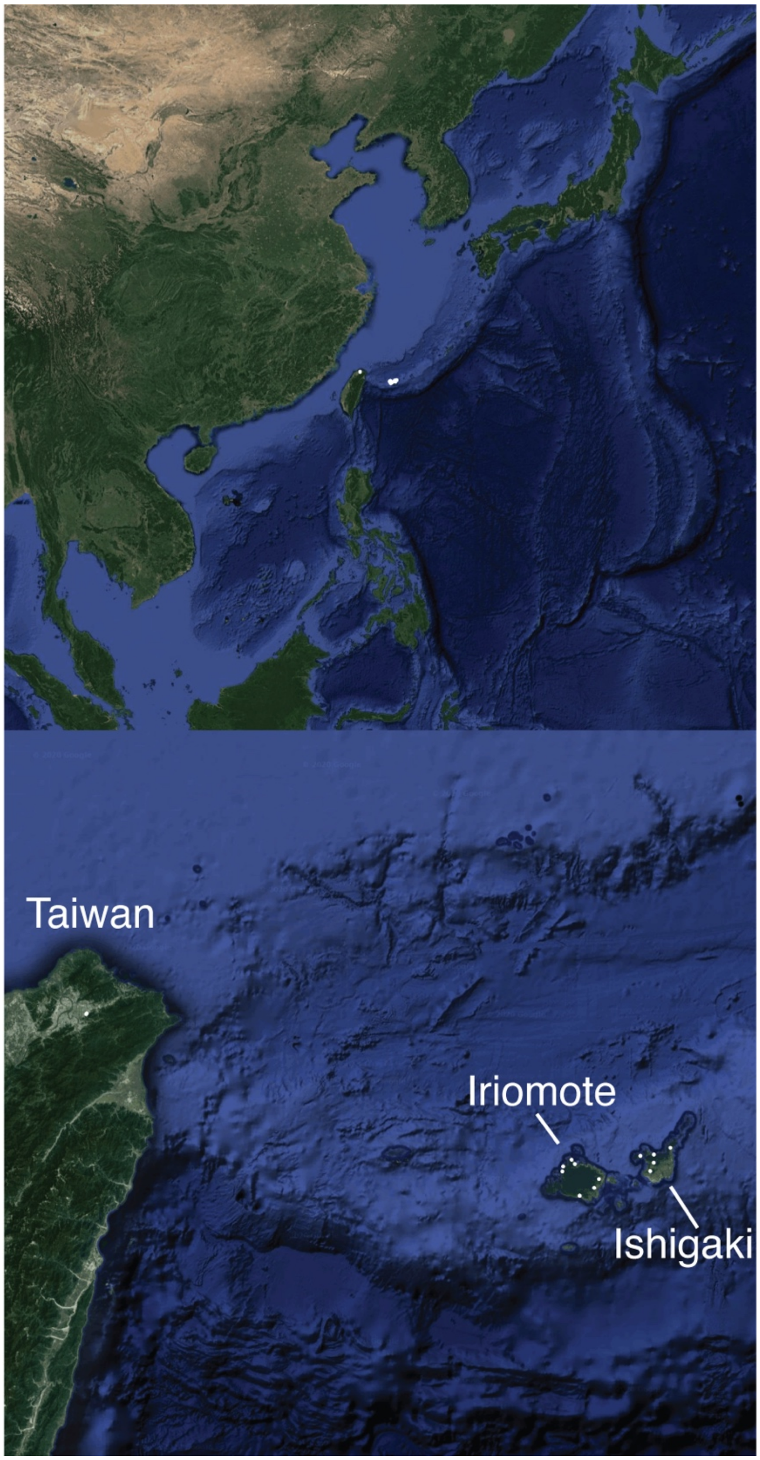


Supplemental Figure 1. Localities of *C. inopinata* strains isolated from Okinawa (islands of Iriomote and Ishigaki) and Taiwan.

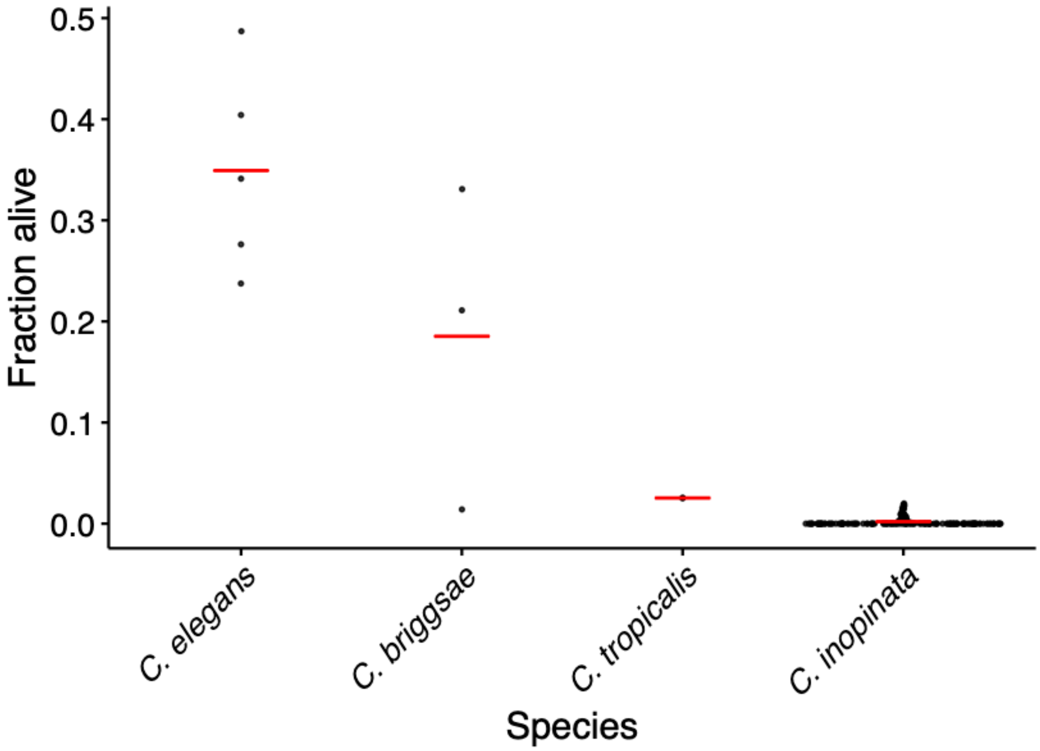


Supplemental Figure 2. *C. inopinata* produces few dauer larvae compared to its close relatives. Strip charts reveal the fraction of starved animals that survive SDS exposure. Red horizontal lines denote means.

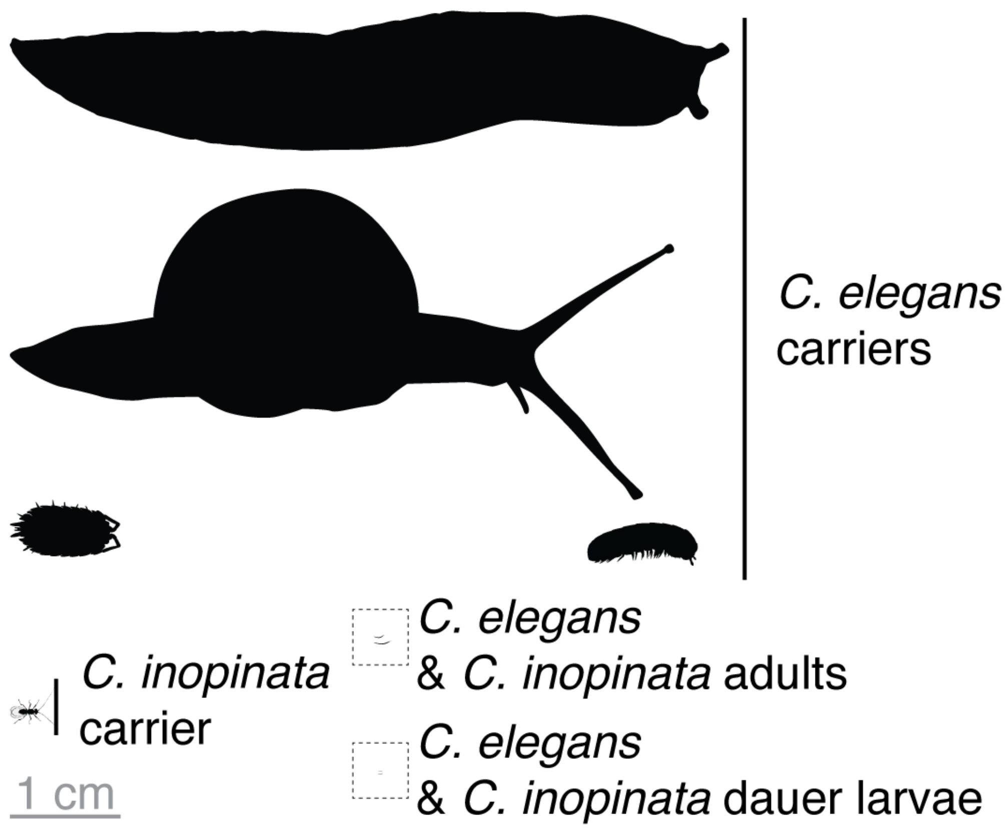


Supplemental Figure 3. Variation in the size of *Caenorhabditis* vectors. As Figure 4, but here including nematodes to scale.
